# The autism and schizophrenia-associated protein CYFIP1 regulates bilateral brain connectivity

**DOI:** 10.1101/477174

**Authors:** Nuria Domínguez-Iturza, Disha Shah, Anna Vannelli, Adrian C. Lo, Marcelo Armendáriz, Ka Wan Li, Valentina Mercaldo, Massimo Trusel, Denise Gastaldo, Manuel Mameli, Annemie Van der Linden, August B. Smit, Tilmann Achsel, Claudia Bagni

## Abstract

Copy-number variants of the *CYFIP1* gene in humans have been linked to Autism and Schizophrenia, two neuropsychiatric disorders characterized by defects in brain connectivity. CYFIP1 regulates molecular events underlying post-synaptic functions. Here, we show that CYFIP1 plays an important role in brain functional connectivity and callosal functions. In particular, we find that *Cyfip1* heterozygous mice have reduced brain functional connectivity and defects in white matter architecture, typically relating to phenotypes found in patients with Autism, Schizophrenia and other neuropsychiatric disorders. In addition, *Cyfip1* deficient mice present deficits in the callosal axons, namely reduced myelination, altered pre-synaptic function, and impaired bilateral-connectivity related behavior. Altogether, our results show that *Cyfip1* haploinsufficiency compromises brain connectivity and function, which might explain its genetic association to neuropsychiatric disorders.

---

The corpus callosum (CC) is the largest axonal commissure in the human brain and connects both cerebral hemispheres. Appropriate callosal microstructure and function are required for bilateral functional connectivity^1,2^. Agenesis of the corpus callosum (ACC), callosotomy or defects in callosal integrity have been linked with deficits in social communication^3^, social behavior^4^, motor coordination and learning^5–7^, and cognitive performance^8,9^. Similarly, corpus callosum abnormalities have been found in patients with diverse neuropsychiatric disorders such as Autism Spectrum Disorders (ASD), Schizophrenia (SCZ), Fragile X syndrome (FXS), attention-deficit hyperactivity disorder (ADHD), Prader-Willi syndrome and 22q11.2 deletion syndrome^3,10–13^. Interestingly, over the last decade, magnetic resonance imaging (MRI) studies in patients with ASD and SCZ have identified functional connectivity and microstructural defects in the white matter as a hallmark of these disorders. Resting state functional MRI (rsfMRI), a measure of brain functional connectivity (FC), is altered in ASD and SCZ patients, showing predominantly reduced long-range functional connectivity^14–19^. Additionally, diffusion tensor imaging (DTI), a measure of axonal integrity and connectivity, is also affected in patients with ASD and SCZ^20–23^. In particular, DTI revealed abnormalities in the corpus callosum of these patients^21,24^. Furthermore, reduced myelination has also been observed in patients with ASD and SCZ^25,26^.

Recently, copy-number variations (CNV) of the chromosomal region 15q11.2 have been associated with the development of neuropsychiatric disorders, especially ASD and SCZ. The proximal region of the long arm of chromosome 15 contains several breakpoints, resulting in different rearrangements^27^. The smallest region that has been linked to neuropsychiatric disorders is contained between the breakpoints BP1 and BP2, i.e., the 15q11.2 locus. CNVs of this region are quite frequent (~1% of the population^28^) and cause a small but significant increase in the risk of developing ASD or SCZ^29,30^. 15q11.2 deletion and duplication carriers show abnormal volume of the corpus callosum among other cerebral abnormalities^31–33^. The chromosomal region contained in the BP1-BP2 encodes four genes: *TUBGCP5*, *NIPA1*, *NIPA2* and *CYFIP1*. Rare CNVs, single nucleotide polymorphisms (SNPs) and point mutations single out *CYFIP1* as the gene most likely causally associated with SCZ^34–36^ and ASD^37,38^. CYFIP1, the Cytoplasmic FMRP interacting protein 1, first described as “specifically Rac1-associated” (Sra-1) protein^39^, is part of the Wave Regulatory Complex (WRC), a heteropentamer formed by WAVE1/2/3, CYFIP1, ABI1/2, NCKAP1 and HPSC300^40,41^. The WRC functions as an Arp2/3 regulator, promoting its actin-nucleating activity, and therefore controlling actin assembly^40^. This process can be activated by Rac1 signaling, which promotes a conformational change in CYFIP1^42^ and its binding to the WRC^41^. In its alternative conformation, CYFIP1 binds FMRP and the cap-binding protein eIF4E, repressing protein translation of specific mRNAs^43^. Both processes, actin dynamics and regulated protein synthesis, are crucial for synaptic development and brain functioning, strengthening the hypothesis that alterations in the *CYFIP1* gene can account for the clinical features of patients with ASD and SCZ.

Deletion of the *Cyfip1* gene in mice leads to embryonic lethality^41,44,45^ while heterozygous mice show aberrant behavior^44^, decreased dendritic complexity, increased number of immature spines^41,45^, as well as electrophysiological defects, such as enhanced mGluR-dependent LTD^44^ and vesicle release probability^46^, suggesting that *Cyfip1* heterozygous mice are a suitable model for the neurological defects observed in BP1-BP2 haploinsufficient patients. Recently, it was shown in zebrafish that CYFIP1 regulates axonal growth of retinal ganglion cells (RGSs)^47^. The role of *Cyfip1* in brain connectivity, however, remains not explored. Here, we show that *Cyfip1* heterozygous mice (*Cyfip1^+/-^*) have reduced functional connectivity and white-matter architecture defects. *Cyfip1* haploinsufficiency leads to a decreased transmission of callosal synapses, reduced myelination in the corpus callosum, and affects motor coordination. Together, our findings suggest that *CYFIP1* is likely the key gene that accounts for the functional connectivity and callosal defects observed in patients with the 15q11.2 deletion, and in other forms of neuropsychiatric disorders with reduced CYFIP1 levels and defects in functional connectivity.

## RESULTS

### Bilateral functional connectivity is impaired in *Cyfip1*^+/-^ mice

Many neuropsychiatric disorders are characterized by impaired brain connectivity. To investigate whether and how *Cyfip1* haploinsufficiency could affect brain networks *in vivo*, we analyzed functional connectivity (FC) in several brain regions using resting-state functional magnetic resonance imaging (rsfMRI). *Cyfip1*^+/-^ adult mice (postnatal day 60, P60) showed decreased functional connectivity (FC) compared to *wild-type* (WT) (compare the lower left and upper right halves of the matrix, Figure 1a). Brain areas that present the most significant differences between WT and *Cyfip1*^+/-^ mice are the cingulate cortex (Cg) and the thalamus (T) (Figure 1b). FC networks as measured by resting state fMRI are characterized by midline symmetry, and correlations between homotopic regions are particularly strong^48,49^. To analyze bilateral FC, seed-based analysis was performed for some of the areas (Figure 1c). Remarkably, FC with the contralateral side of the seed region was particularly affected in the hippocampus, somatosensory and motor cortices (Figure 1d). In conclusion, our results show significant defects in functional connectivity, especially of the bilateral connections.

**Figure 1.**
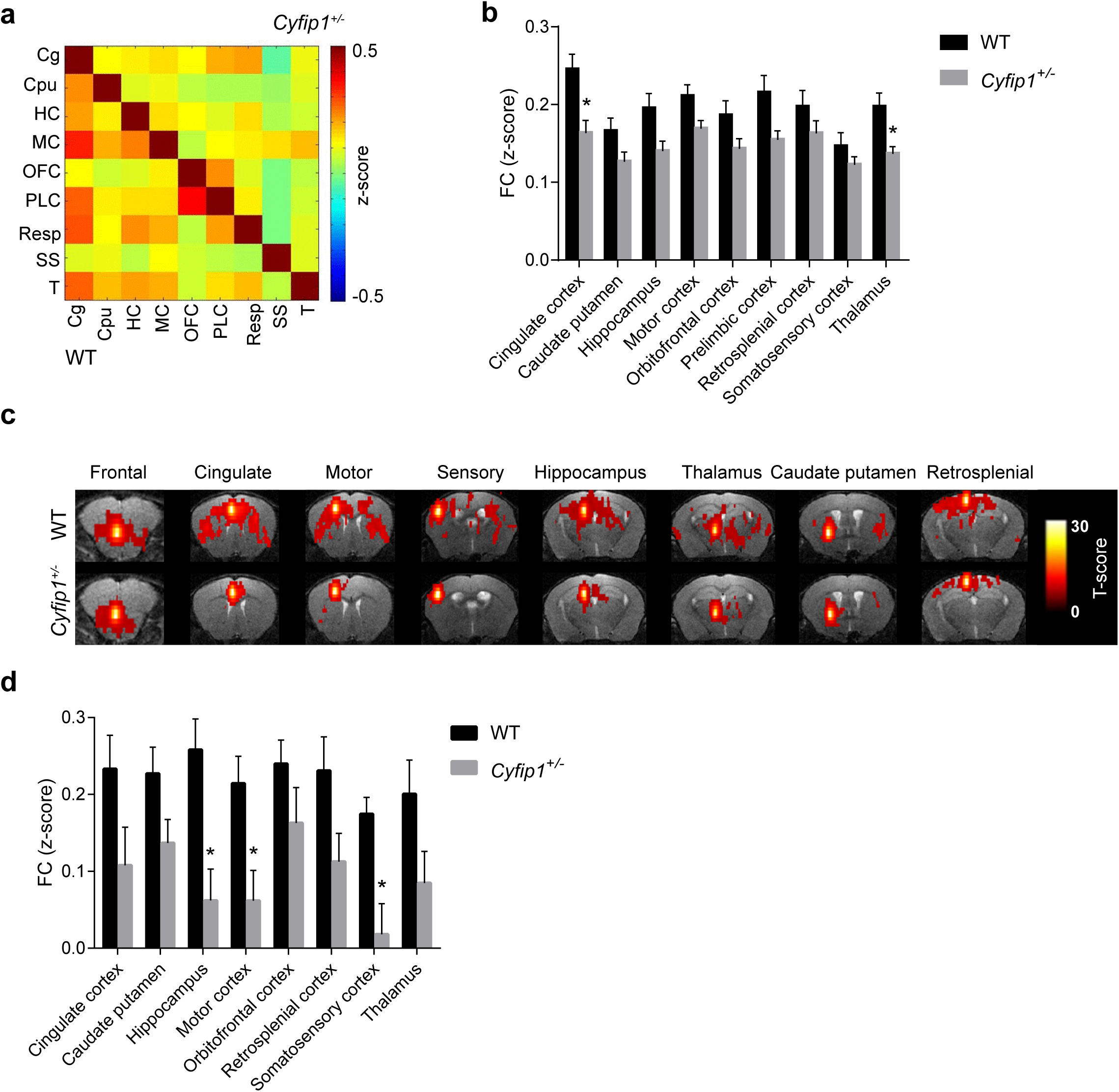
*Cyfip1*^+/-^ mice show reduced functional connectivity. (a) Functional connectivity matrices of WT and *Cyfip1*^+/-^ mice (lower and upper half of the matrix, respectively) (postnatal day 60, P60), in which functional correlation (z-score) between pairs of regions is represented by a color scale (abbreviations: Cg=cingulate cortex, Cpu=caudate putamen, HC=hippocampus, MC=motor cortex, OFC=orbitofrontal cortex, PLC=prelimbic cortex, Resp=retrosplenial cortex, SS=somatosensory cortex, T=thalamus). (b) Based on the matrix in (a), average functional connectivity strength of each region with all other regions is plotted for WT and *Cyfip1*^+/-^ mice (WT n=15 and *Cyfip1*^+/-^ n=17 mice) (mean ± SEM; Holm-Sidak t-test; Cg, p=0.0178 and T, p=0.0196). (c) Seed-based analysis represented by functional connectivity maps for WT and *Cyfip1*^+/-^ animals. The strength of connectivity for the seed region indicated above each image is mapped by a color scale representing the *T*-score. (d) Average bilateral functional connectivity strength for selected brain regions in WT and *Cyfip1*^+/-^ mice (WT n=15 and *Cyfip1*^+/-^ n=17 mice) (mean ± SEM; Holm-Sidak t-test; MC p=0.0452, SS p=0.0157, HC p=0.0148).

### Aberrant white matter architecture in *Cyfip1*^+/-^ mice

Bilateral connectivity mostly passes through the corpus callosum and abnormalities in this brain structure are a hallmark of neuropsychiatric disorders. To determine whether the observed functional deficits are due to structural axonal abnormalities in the corpus callosum, we performed diffusion tensor imaging (DTI)^50^ in WT and *Cyfip1*^+/-^ animals. DTI is one of the most widely used techniques to analyze white matter architecture and integrity. DTI yields several variables, of which the fractional anisotropy (FA) is the most widely used. High FA values are indicative of fiber-like structures and therefore, FA provides an estimate of axonal architecture^50^. In Figure 2a, the FA values across WT and *Cyfip1*^+/-^ brains were color-coded, with higher values in red highlighting white matter tracts such as the CC (prominent in the first to fifth image). FA was generally reduced throughout the brain in the *Cyfip1*^+/-^ mice (compare the upper and lower rows); except for the fimbria that showed slightly increased FA. The differential map of WT and *Cyfip1*^+/-^ FA values showed that the highest reduction in FA was detected in the CC, in particular in the genu, its anterior part. (Figure 2b; first image). We therefore calculated the DTI parameters in the corpus callosum. No statistical differences were observed in the mean (MD), axial (AD), or radial diffusivity (RD), indicating that there are no gross defects in the white matter structure. In contrast, fractional anisotropy (FA) was significantly lower in the corpus callosum of *Cyfip1*^+/-^ mice (Figure 2c), which might indicate changes in axonal thickness or myelination.

**Figure 2.**
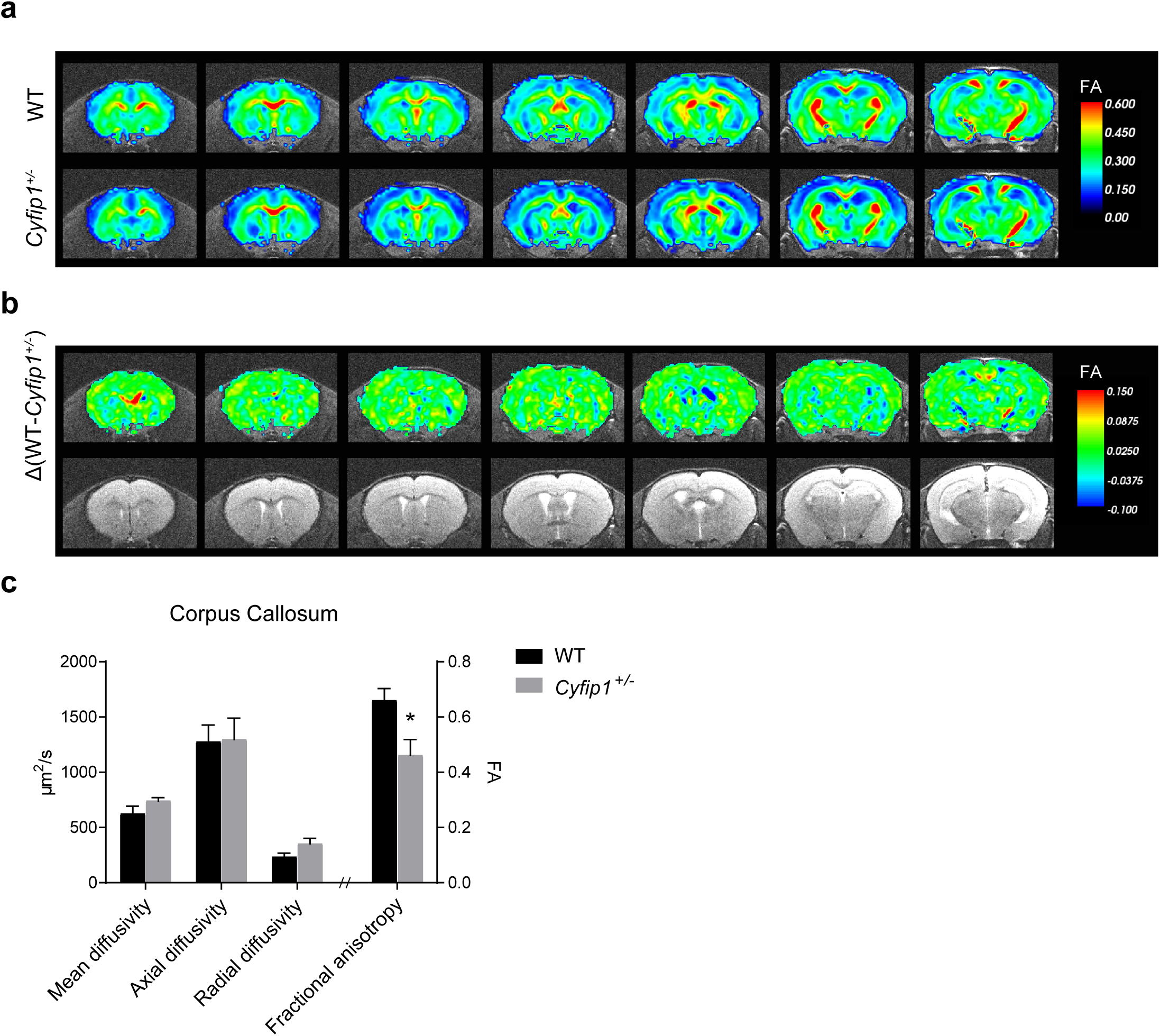
*Cyfip1*^+/-^ mice show defects in callosal architecture. (a) Diffusion Tensor Imaging (DTI) of WT and *Cyfip1*^+/-^ mice at postnatal day 60 (P60). Fractional anisotropy (FA) maps in which average FA values for WT and *Cyfip1*^+/-^ mice are represented by a color scale (WT n=7 and *Cyfip1*^+/-^ n=6 mice). (b) Upper, FA differential maps between WT and *Cyfip1*^+/-^ (Δ(WT-*Cyfip1*^+/-^)). Lower, representative anatomical MRI images as reference. (c) The graph shows mean diffusivity (MD), axial diffusivity (AD), radial diffusivity (RD) and fractional anisotropy (FA) in the corpus callosum (WT n=7 and *Cyfip1*^+/-^ n=6 mice) (mean ± SEM; t-test; FA p=0.0165).

To determine which of the two properties is affected, we performed electron microscopy of the corpus callosum (CC). Representative micrographs from the anterior part of the CC were selected (Figure 3a), and myelinated axons were automatically identified and parameterized. For each axon, the diameter and the myelin thickness were determined. In addition, the g-ratio, i.e., the ratio of the internal over the external axon diameter (Figure 3b) was calculated. We observed no changes in the axonal diameter between the two genotypes, whereas the myelin thickness was reduced (Figure 3c). Consequently, the g-ratio increased across all axon sizes (Figure 3c). In conclusion, *Cyfip1* haploinsufficiency causes defects in functional connectivity, especially of the callosal projections that connect the two cortical hemispheres, which correlates with a reduced myelin thickness of the callosal fibers.

**Figure 3.**
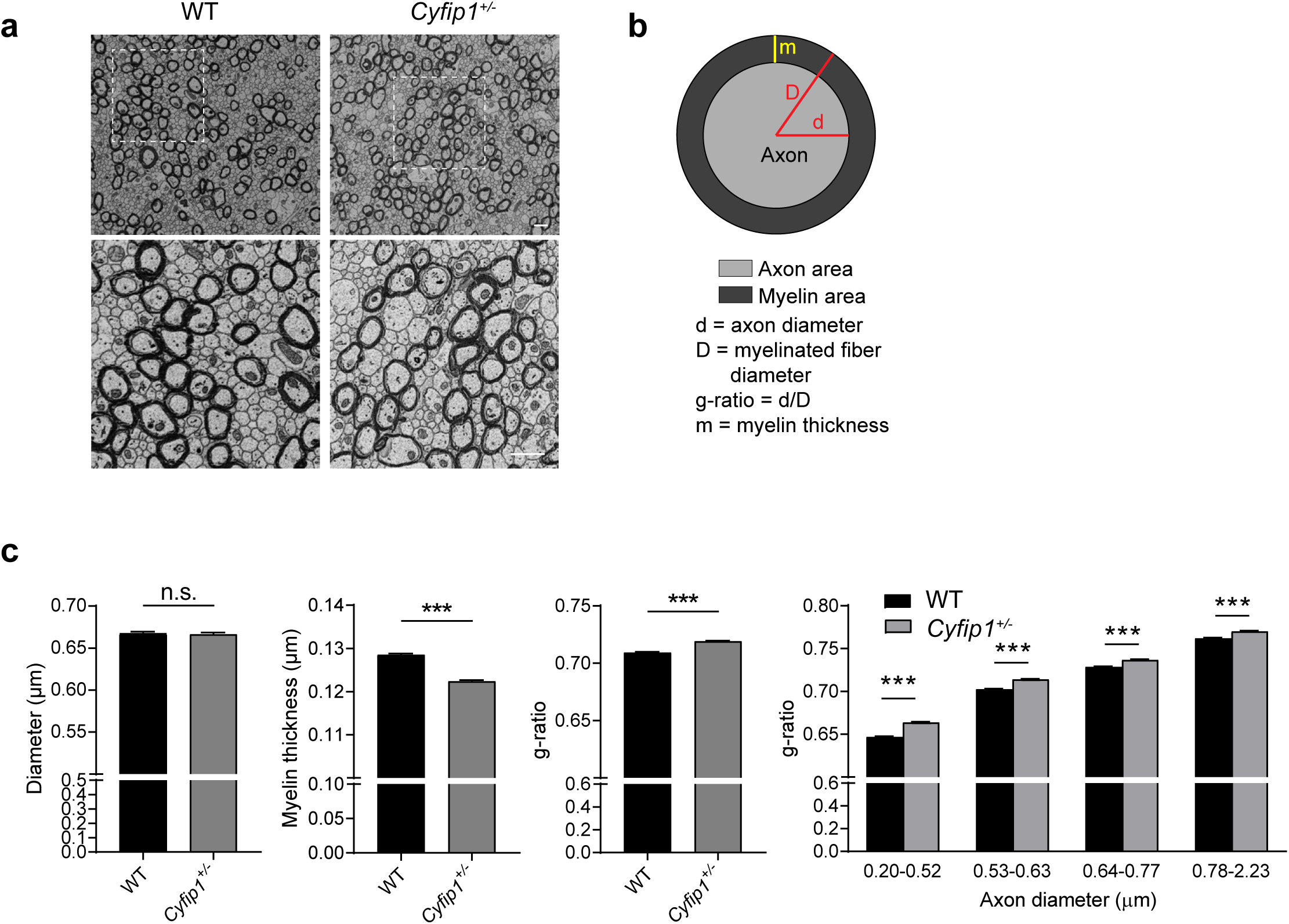
*Cyfip1*^+/-^ mice show defects in callosal myelination. (a) Representative electron micrographs of axons in the genu of the corpus callosum (CC) in WT and *Cyfip1*^+/-^ mice at P60. Lower panel, zoom image (scale bar 1 µm). (b) Schematic of the measured parameters. (c) Graphs show average axonal diameter, myelin thickness and g-ratio. The histograms on the right show the g-ratio of axons with different diameters (n >10000 axons for each genotype, n=3 mice for each genotype, mean ± SEM; Mann-Whitney test; myelin thickness p<0.0001, g-ratio p<0.0001; Two-way ANOVA with Holm-Sidak’s multiple comparison test, g-ratio p<0.0001).

### *Cyfip1* haploinsufficiency leads to altered callosal presynaptic function

It is well established that neuronal activity can regulate callosal myelination^51^. To investigate whether *Cyfip1* deficiency leads to defects in neuronal activity, we measured spontaneous activity in the somatosensory cortex of acute brain slices using microelectrode arrays (MEAs) (Figure 4a). We found that both the spike and burst rate were significantly reduced in the *Cyfip1*^+/-^ cortices (Figure 4b-d), indicating that there are perceptible network alterations in the adult brain. This reduction in spontaneous activity could explain the reduced myelin thickness of callosal axons, which would in turn affect their activity. To study the latter, we extracellularly stimulated the corpus callosum to elicit EPSCs in pyramidal cells of the layer II/III^52^, where contralateral projections arrive through the callosal tract (Figure 5a). Upon challenging callosal axons with trains of stimulation, slices from wild type mice presented short-term synaptic depression. In contrast, in *Cyfip1*^+/-^ slices we observed the absence of paired-pulse depression (Figure 5b and c), indicating altered presynaptic function (see Discussion). Altogether, this evidence corroborates the hypothesis that in *Cyfip1*^+/-^ mice, bilateral transmission via callosal tracts is impaired.

**Figure 4.**
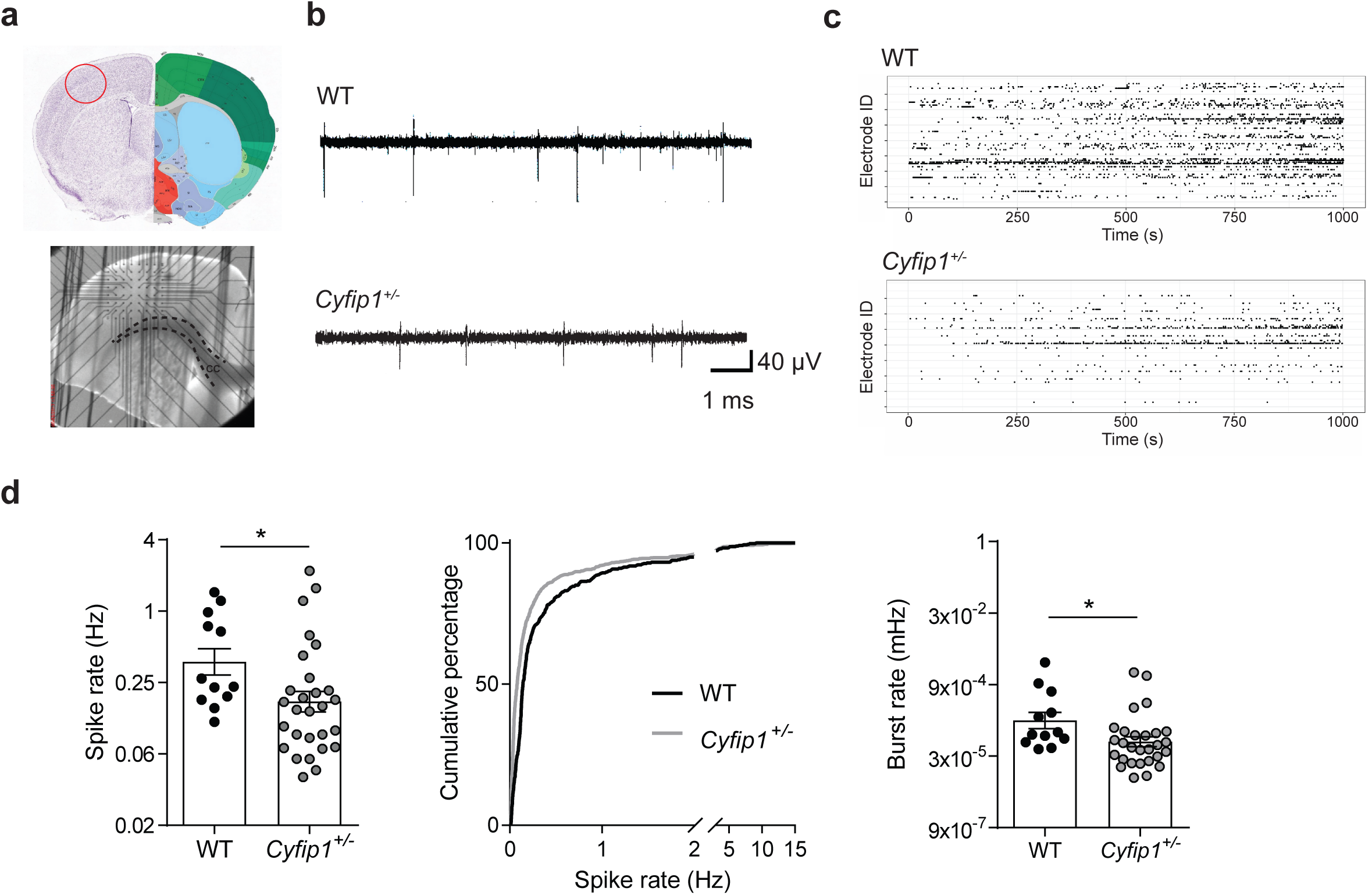
Spontaneous activity is reduced in *Cyfip1*^+/-^ adult mice. (a) Upper panel, representative image of a P60 brain slice (Allen Brain Atlas, brain regions are color-contrasted) where the recorded area is highlighted by a red circle. Lower panel, representative image of the P60 somatosensory cortex slice positioned on the Microelectrode Array (MEA) system. Dashed black lines delineate the corpus callosum (CC). The field of electrodes is visible above the CC. (b) Representative traces of single electrodes recording from WT and *Cyfip1*^+/-^ slices. (c) Representative raster plots of all 59 electrodes showing the spike events in WT and *Cyfip1*^+/-^ cortical slice recordings. The x-axis corresponds to the recording time and the y-axis to the electrode ID. (d) Quantification of the spike rate (left) and burst rate (right, both on a logarithmic scale) in WT and *Cyfip1*^+/-^ brain slices at P60 (WT n=12 slices, 6 mice, and *Cyfip1*^+/-^ n=28 slices, 13 mice) (mean ± SEM; two sample t-test; p=0.032 for spike rate and p=0.027 for burst rate). Center, cumulative frequency distribution of the spike rate for WT and *Cyfip1*^+/-^ mice (Kolmogorov-Smirnov test, p<0.001).

**Figure 5.**
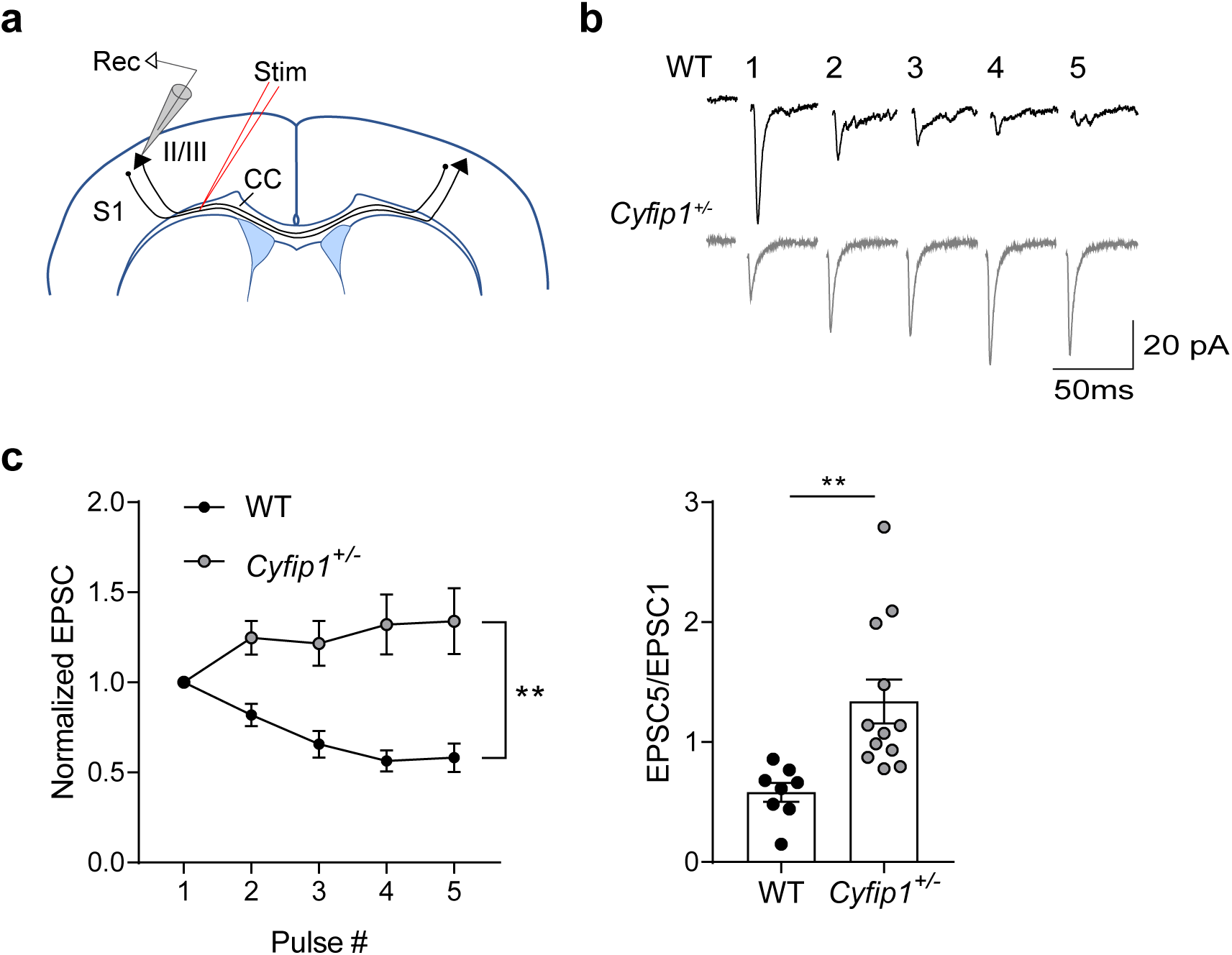
Reduced presynaptic function in *Cyfip1*^+/-^ adult mice. (a) Schematic of the experimental setup. (b) Representative sample traces induced in layers II/III cortical neurons at P60 by trains of callosal stimulation (5 pulses, 20 Hz). Recordings obtained from WT (black) and *Cyfip1*^+/-^ (gray) (c) Left, graph of the normalized amplitude of EPSCs, and right, histogram and scatter plot of the amplitudes of the fifth EPSCs, relative to the first. (WT n=8 cells, 3 mice and *Cyfip1*^+/-^ n=12 cells, 3 mice) (mean ± SEM; Two-way Repeated Measures ANOVA, genotype main effect p=0.0017, and interaction p<0.0001; t-test p=0.0046).

### *Cyfip1*^+/-^ mice show motor coordination defects

Callosal abnormalities have been associated with motor deficits, particularly with motor coordination, both in humans^6^ and in mice^53,54^. We therefore used the accelerating rotarod test (Figure 6a) to investigate how the impaired bilateral connectivity and callosal transmission in *Cyfip1*^+/-^ mice affect motor function. As shown in Figure 6b-c, C*yfip1^+/-^* mice fall off significantly earlier than their WT littermates, indicating reduced motor coordination. Of note, this defect was ameliorated after repeated trials (Figure 6c), suggesting intact motor learning. To exclude that the abnormal coordination was due to reduced locomotion, general activity of *Cyfip1*^+/-^ mice was measured in the open field. *Cyfip1*^+/-^ animals moved normally (Figure 6d-f) indicating that locomotion was normal and that the defect on the rotarod is indeed due to impaired motor coordination. In sum, we gathered compelling electrophysiological, anatomical, functional and behavioral evidence, which all demonstrate defects in callosal connections of the adult *Cyfip1*^+/-^ mouse brain.

**Figure 6.**
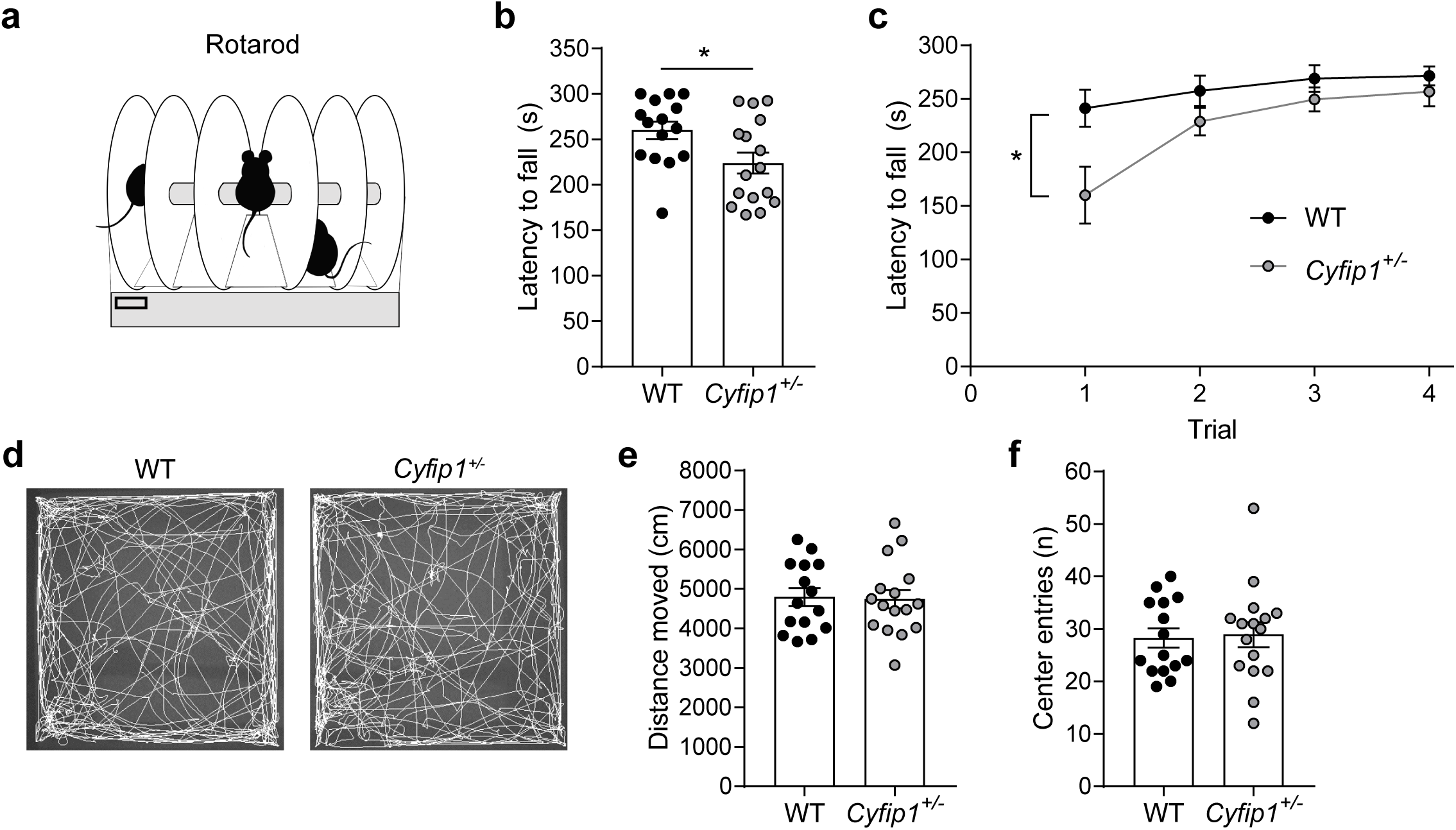
Motor coordination is reduced in *Cyfip1*^+/-^ mice. (a) Schematic of the rotarod test. (b) Average latency to fall measured in the accelerating rotarod (WT n=15 and *Cyfip1*^+/-^ n=16 mice) (mean ± SEM; two-tailed t-test; p=0.024). (c) Latency to fall per trial measured in the accelerating rotarod (WT n=15 and *Cyfip1*^+/-^ n=16 mice) (mean ± SEM; Two-way Repeated-Measures ANOVA; p=0.024 for genotype). (d) Representative movement tracks of WT and *Cyfip1*^+/-^ mice in the open field. (e and f) Open field parameters measured for WT and *Cyfip1*^+/-^ mice; distance moved and center entries, respectively (WT n=15 and *Cyfip1*^+/-^ n=16 mice) (mean ± SEM; two-tailed t-test).

## DISCUSSION

Here we describe an impairment in callosal architecture and transmission in a mouse model of *Cyfip1* haploinsufficiency. These callosal defects have an impact on functional connectivity and on motor coordination which might explain the genetic association of CYFIP1 with neuropsychiatric disorders.

We observed impaired bilateral functional connectivity, which correlates with several kinds of alterations in the collosal axons: in the microstructure as determined by DTI, which correlates with reduced myelin thickness (Figure 2 and 3), and in pre-synaptic transmission (Figure 5). Regarding pre-synaptic function, we observe that trains of stimulation lead to short-term synaptic depression in WT but not *Cyfip1*^+/-^ slices (Figure 5). This could be caused by changes in neurotransmitter release probability^55^, which would be higher in WT slices, but other presynaptic alterations could also be the cause. Arguably, changes in synaptic transmission have an impact on spontaneous activity, as we observed (Figure 4), and neuronal activity modulates myelination of the callosal axons^51^. In this model, the primary defect would lie in the callosal axons. On the other hand, reduced myelination affects signal propagation along the axons and could therefore be the primary cause of the callosal defects. Further studies will be needed to elucidate this cause-consequence relation.

Even though the largest reduction in FA in the *Cyfip1*^+/-^ mice was observed in the CC, other axonal tracts may also be affected. CYFIP1 is expressed across the whole brain^45,56^ and therefore axonal defects in the *Cyfip1*^+/-^ mice may be generalized. Indeed, both fractional anisotropy and functional connectivity are generally reduced across the whole brain in *Cyfip1*^+/-^ mice. Additionally, defects in FC and white matter architecture have been reported across the brain in patients with ASD or SCZ^15,19,22,57^. Long-range connectivity problems between brain areas may also contribute, in addition to the callosal defects, to the observed deficits in motor coordination.

Importantly, patients with the 15q11.2 BP1–BP2 microdeletion syndrome present callosal defects. Notably, a high proportion of 15q11.2 patients (42%) show motor delay^28^, which may correlate with the reduced motor coordination we observed (Figure 6). In addition, motor coordination deficits have been observed in patients with ASD^58,59^. Of note, the BP1–BP2 region contains four genes. Here, we show that the callosal defects are caused by the sole deletion of the *Cyfip1* gene, giving further evidence that human *CYFIP1* is the causative gene in the BP1–BP2 region.

Alterations in the callosal region are not only found in patients with 15q11.2 BP1-BP2 deletions and duplications^31–33^, they are also a hallmark of ASD^24^ and SCZ^21^. Thus, variations in other chromosomal regions genetically associated with ASD or SCZ present similar defects in brain structure and connectivity as the *Cyfip1* haploinsufficient mice. For instance, 22q11.2 deletion carriers, a chromosomal rearrangement that is associated with SCZ, present structural abnormalities in several brain regions^60,61^ which are recapitulated in a mouse model for the disease (Df(16)A^+/-^)^62^. Similarly, studies in humans carrying the autism-associated 16p11.2 microdeletion showed reduced prefrontal connectivity and microstructural defects in the corpus callosum. These defects were also found in the mouse model for the disease, in which the callosal abnormalities were associated with increased axonal g-ratio^63^. Finally, genetic variations in the CNTNAP2 gene have been associated with ASD^64^. Absence of CNTNAP2 leads to defects in callosal transmission and cortical myelination^52^, as well as reduced functional connectivity and aberrant g-ratio of the callosal axons^65,66^. Taken together, these studies point out that neuropsychiatric disorders of different genetic origins present similar defects in brain connectivity and white matter architecture, especially in the corpus callosum, highlighting the importance of brain connectivity as a possible point of convergence for several neuropsychiatric disorders^67^. In addition, some of the areas that show reduced functional connectivity in the *Cyfip1*^+/-^ mice are part of the default mode network (DMN), a brain network that reflects functional connectivity across different brain regions during conscious-inactive tasks^68,69^. Although the exact function of the DMN is still unknown, DMN defects have been found in patients with SCZ and ASDs^70–73^ and have been suggested to be key for understanding the pathophysiology of these disorders^71,74^. All together, the connectivity deficits caused by CYFIP1 deficiency could therefore have a major contribution in the pathogenesis of neuropsychiatric disorders.

In conclusion, here we found that *Cyfip1* haploinsufficiency is directly responsible for key phenotypes of patients with ASD and SCZ, namely the alterations in anatomic and functional connectivity, thereby explaining the increased incidence of these pathologies in patients with 15q11.2 deletions. Importantly, this pathological mechanism might not be limited to neuropsychiatric disorders caused by CNVs in the 15q11.2 region. CYFIP1 has been found dysregulated at the protein level in SCZ independently of mutations in the CYFIP1 locus^75^, indicating that other genetic and/or environmental factors may converge on the regulation of CYFIP1 abundance or activity, rendering the protein a hub for the development of SCZ and perhaps also related disorders.

## METHODS

### Animal care

Animal housing and care was conducted according to the institutional guidelines that are in compliance with national and international laws and policies (Belgian Royal Decree of 29 May 2013, European Directive 2010/63/EU on the protection of animals used for scientific purposes of 20 October 2010 and Swiss Loi fédérale sur la protection des animaux 455). All the experimental procedures performed in Switzerland complied with the Swiss National Institutional Guidelines on Animal Experimentation and were approved by the Cantonal Veterinary Office Committee for Animal Experimentation. In all cases special attention was given to the implementation of the 3 R’s, housing and environmental conditions and analgesia to improve the animals’ welfare.

A 12h light/dark cycle was used, and food and water were available *ad libitum*. The *Cyfip1*^+/-^ mouse line, generated by gene trap at the Sanger Institute, UK, was kindly provided by Seth G.N. Grant. The gene trap cassette was inserted between exon 12 and 13. Molecular characterization of the *Cyfip1*^+/-^ mice was previously done by De Rubeis et al. 2013^41^. C57/Bl6 wild type (WT) and *Cyfip1* heterozygous (*Cyfip1*^+/-^) male mice at postnatal day 60-70 (P60-70) were used. To generate the experimental animals, *Cyfip1*^+/-^ male were crossed with WT C57Bl6 females. Animal manipulation was performed according to the ECD protocol approved by the Institutional ethical committee of the KU Leuven and Cantonal Veterinary Office Committee for Animal Experimentation.

### Resting state functional magnetic resonant imaging and diffusion tensor imaging

#### Animals

For the MRI handling procedures, adult male mice were anesthetized with 2.5% isoflurane (IsoFlo, Abbott, Illinois, USA), which was administered in a mixture of 70% nitrogen (400 cc/min) and 30% oxygen (200 cc/min).

RsfMRI imaging procedures were performed as previously described^76,77^. In brief, a combination of medetomidine (Domitor, Pfizer, Karlsruhe, Germany) and isoflurane was used to sedate the animals. After positioning of the animal in the scanner, medetomidine was administered subcutaneously as a bolus injection (0.3 mg/kg), after which the isoflurane level was immediately decreased to 1%. Five minutes before the rsfMRI acquisition, isoflurane was decreased to 0.4%. RsfMRI scans were consistently acquired 40 min after the bolus injection, during which the isoflurane level was maintained at 0.4%. After the imaging procedures, the effects of medetomidine were counteracted by subcutaneously injecting 0.1 mg/kg atipamezole (Antisedan, Pfizer, Karlsruhe, Germany).

Regarding DTI measurements, after the handling procedures under isoflurane (2.5%), isoflurane levels were decreased to 1.5% and maintained throughout the scanning procedure. The physiological status of all animals was monitored throughout the imaging procedure. A pressure sensitive pad (MR-compatible Small Animal Monitoring and Gating system, SA Instruments, Inc.) was used to monitor breathing rate and a rectal thermistor with feedback controlled warm air circuitry (MR-compatible Small Animal Heating System, SA Instruments, Inc.) was used to maintain body temperature at (37.0 ± 0.5) °C

#### Imaging procedures

RsfMRI procedures were performed on a 9.4T Biospec MRI system (Bruker BioSpin, Germany) with the Paravision 5.1 software (www.bruker.com). Images were acquired using a standard Bruker cross coil set-up with a quadrature volume coil and a quadrature surface coil for mice. Three orthogonal multi-slice Turbo RARE T2-weighted images were acquired to render slice-positioning uniform (repetition time 2000 ms, effective echo time 33 ms, 16 slices of 0.5 mm). Field maps were acquired for each animal to assess field homogeneity, followed by local shimming, which corrects for the measured inhomogeneity in a rectangular VOI within the brain. Resting-state signals were measured using a T2*-weighted single shot EPI sequence (repetition time 2000 ms, echo time 15 ms, 16 slices of 0.5 mm, 150 repetitions). The field-of-view was (20 × 20) mm² and the matrix size (128 × 64).

DTI was performed on a 7T Pharmascan system (Bruker BioSpin, Germany). Images were acquired using a Bruker cross coil set-up with a transmit quadrature volume coil and a receive-only surface array for mice. Three orthogonal multi-slice Turbo RARE T2-weighted images were acquired to render slice-positioning uniform (repetition time 2500 ms, effective echo time 33 ms, 18 slices of 0.5 mm). Field maps were acquired for each animal to assess field homogeneity, followed by local shimming, which corrects for the measured inhomogeneity in a rectangular VOI within the brain. DTI images were acquired using a multislice two-shot spin-echo EPI sequence (repetition time 5500 ms, echo time 23.23 ms, 18 slices of 0.5 mm, b=800 s/mm², 60 DW direction). The field of view was (20 x 20) mm² and the matrix size (96 x96).

#### Image processing

Pre-processing of the rsfMRI and DTI data, including realignment, normalization and smoothing (for the rsfMRI data), was performed using SPM8 software (Statistical Parametric Mapping, http://www.fil.ion.ucl.ac.uk). First, all images within each session were realigned to the first image. This was done using a least-squares approach and a 6-parameter (rigid body) spatial transformation. For the analyses of the rsfMRI data, motion parameters resulting from the realignment were included as covariates to correct for possible movement that occurred during the scanning procedure. Second, all datasets were normalized to a study-specific EPI template. The normalization steps consisted of a global 12-parameter affine transformation followed by the estimation of the nonlinear deformations. Finally, in plane smoothing was done for the rsfMRI data using a Gaussian kernel with full width at half maximum of twice the voxel size (0.31 × 0.62 mm²). All rsfMRI data were filtered between 0.01-0.1 Hz using the REST toolbox (REST1.7, http://resting-fmri.sourceforge.net).

Regions-of-interest (ROIs) were delineated using the MRicron software (MRicron version 6.6, 2013, http://www.mccauslandcenter.sc.edu/mricro/): cingulate cortex, caudate putamen, hippocampus, motor cortex, orbitofrontal cortex, prelimbic cortex, retrosplenial cortex, somatosensory cortex and thalamus. The rsfMRI data were first analyzed using a region-of-interest (ROI) correlation analysis, where pairwise correlation coefficients between each pair of ROIs were calculated and z-transformed using an in-house program developed in MATLAB (MATLAB R2013a, The MathWorks Inc. Natick, MA, USA). Mean z-transformed FC matrices were calculated for each group. For each brain region mean FC, i.e. mean FC of each region with other regions in the matrix, and bilateral FC was calculated. Statistical analyses of the pairwise correlations included two-way ANOVAs to compare groups and brain regions using the Holm-Sidak correction for multiple comparisons.

Next, a seed-based analysis was performed identifying all functional connections of the specific region to other voxels in the brain. Seed-based analyses were performed by first computing individual z-transformed FC-maps of the respective region using the REST toolbox, after which mean statistical FC-maps were calculated for each group in SPM8.

Each individual DTI FA-map was warped to a reference anatomical image. Then, average FA-maps for WT and *Cyfip1*^+/-^ mice were computed and a differential map (Δ(WT-*Cyfip1*^+/-^)) was calculated. Finally, DTI parameters (i.e. radial diffusivity, axial diffusivity, mean diffusivity and fractional anisotropy) were computed with MATLAB and statistical analyses were performed using two sample t-tests to compare group differences in the corpus callosum.

### Electron microscopy

Adult mice from both genotypes were perfused, via the heart, with 100 ml of a buffered mix of 2.5% glutaraldehyde and 2.0% paraformaldehyde in 0.1 M phosphate buffer (pH 7.4), and then left for 2h before the brain was removed, embedded in 5% agarose, and sagittal vibratome sections cut at 80 µm thickness through the midline. These sections were then post-fixed in potassium ferrocyanide (1.5%) and osmium (2%), then stained with thiocarbohydrazide (1%) followed by osmium tetroxide (2%). They were then stained overnight in uranyl acetate (1%), washed in distilled water at 50°C before being stained with lead aspartate at the same temperature. They were finally dehydrated in increasing concentrations of alcohol and then embedded in spurs resin and hardened at 65°C for 24h between glass slides. The regions containing the corpus callosum were trimmed from the rest of the section using a razor blade and glued to a blank resin block. One micrometer thick sections were then cut from the block face and mounted onto silicon wafers of 1 cm diameter.

The sections were imaged inside a scanning electron microscope (Zeiss Merlin, Zeiss NTS) at a voltage of 2 kV and image pixel size of 7 nm. Backscattered electrons were collected with a Gatan backscattered electron detector with a pixel dwell time of 1 µs. Multiple images of the corpus callosum were collected and tiled using the TrakEM2 plugin^78^ in the FIJI software (www.fiji.sc).

A custom-made MATLAB code was used to identify and parameterize myelinated axons within the EM images. Inner (axon) and outer (axon + myelin) areas were segmented and their corresponding circular equivalent diameters calculated. The g-ratio was calculated as the ratio between the internal and external diameter (d/D) (Figure 3b).

### Multi Electrode Array (MEA) recordings

Cortical slices were prepared from adult male mice to study the effects of *Cyfip1* haploinsufficiency in spontaneous activity. Briefly, the mice were anesthetized with isoflurane and the brains were quickly removed and placed in bubbled ice-cold 95% O_2_/5% CO_2_-equilibrated solution containing (in mM): NaCl 125, KCl 2.5, NaH_2_PO_4_ 1, MgCl_2_ 1.2, CaCl_2_ 0.6, Glucose 11, NaHCO_3_ 26 (95% O_2_, 5% CO_2_). Coronal brain slices (350 µm thick) were made using a vibratome (Leica Biosystem). Slices were kept for 30 minutes at 33°C and for 1 hour at RT in ACSF containing (in mM): NaCl 125, KCl 2.5, NaH_2_PO_4_ 1, MgCl_2_ 2, CaCl_2_ 1, Glucose 11, NaHCO_3_ 26 (95% O_2_, 5% CO_2_).

A single slice was placed in a MEA chamber (Multichannel system, Reutlingen, Germany) positioning the electrode array on the somatosensory cortex. Each MEA is composed of 59 TiN electrodes organized in a 8×8 grid layout. The electrode distance is 200 µm, and the electrode diameter is 30 µm. The slice was stabilized in the MEA chamber using a platinum anchor and was continuously perfused with modified ACSF containing (in mM): MgCl_2_ 0.75, CaCl_2_ 1.4, (95% O_2_, 5% CO_2_). Neuronal activity, sampled at 10 kHz and filtered at 100 Hz, was recorded with a MEA2100-acquisition System (Multichannel system, Reutlingen, Germany). The data was acquired with the MultiChannel Experimenter 2.8 (Multichannel system, Reutlingen, Germany) and extracted for analysis using the MultiChannel Analyzer 2.6 (Multichannel system, Reutlingen, Germany). For spike detection, the threshold was set at −5.5 s.d. of the noise amplitude. An event was considered a burst if it was composed of at least 5 consecutive spikes with an inter-spike interval of less than 50 ms. Data formatting, detection of bursts and events as well as the statistical analyses were performed with a custom-written script in R (software version 3.4.3).

### Patch clamp recordings

Animals were decapitated and the brain was quickly extracted. Slicing was done in bubbled ice-cold 95% O_2_/5% CO_2_-equilibrated solution containing (in mM): choline chloride 110; glucose 25; NaHCO_3_ 25; MgCl_2_ 7; ascorbic acid 11.6; sodium pyruvate 3.1; KCl 2.5; NaH_2_PO_4_ 1.25; CaCl_2_ 0.5. Coronal slices (250 µm) were stored at ∼22°C in 95% O_2_/5% CO_2_-equilibrated artificial cerebrospinal fluid (ACSF) containing (in mM): NaCl 124; NaHCO_3_ 26.2; glucose 11; KCl 2.5; CaCl_2_ 2.5; MgCl_2_ 1.3; NaH_2_PO_4_ 1. Recordings (flow rate of 2.5 ml/min) were made under an Olympus-BX51 microscope (Olympus) at 32°C. Currents were amplified, filtered at 5 kHz and digitized at 20 kHz. Access resistance was monitored by a step of −4 mV (0.1 Hz). Experiments were discarded if the access resistance increased more than 20%. A bipolar stimulating electrode was placed in the corpus callosum, and EPSCs were evoked in layer II/III pyramidal neurons while holding the cells at −60 mV. 5 pulses at 20 Hz were delivered at 0.1 Hz to measure paired pulse ratios. The internal solution contained (in mM): CH_3_CsO_3_S 120; CsCl 10; HEPES 10; EGTA 10; sodium creatine-phosphate 5; Na_2_ATP 4; Na_3_GTP 0.4; pH 7.3

### Behavioral assays

#### Open field

Exploration was observed in an open field setup with a square surface area of 45 cm². Mice were placed in the open field for 10 min and movements were recorded with EthoVision tracking software (Noldus, Wageningen, The Netherlands). A central zone was defined as a square surface area 15 cm equidistantly from all 4 walls of the open field. Total distance traveled as well as center entries were measured.

#### Rotarod

Motor coordination was tested on the accelerating rotarod as previously described (Ugo Basile Model 7500, Gemonio, Italy)^79^. Briefly, mice were first trained for 2 min on the rotarod at constant speed (4 rpm). They were subsequently tested on four 5-min trials interleaved with 10 min rest. During the test trials, mice were placed on a rotating rod that accelerated from 4 to 40 rpm in 5 min and the latency to fall off the rod was recorded.

### Statistics

Statistical analyses were performed with SPSS and Matlab for the MRI and DTI experiments, and with GraphPad Prism or R for EM, electrophysiology, and behavioral experiments. The statistical tests used are listed in the respective Figure legends. In brief, unpaired Student’s t-test or non-parametric Mann–Whitney U-test were used for comparisons between the two groups. Multiple-t-test with Holm-Sidak multiple comparison correction was used for the rsfMRI experiments. Two-way ANOVA with Holm-Sidak’s multiple comparison test was used for analysis of the g-ratio across different axonal sizes. For cumulative frequency distribution, Kolmogorov-Smirnov test was used. Paired pulse ratios and latency to fall were compared with Two-way Repeated Measures ANOVA. For all analysis, P values < 0.05 were considered significant. Results were presented as mean ± standard error of the mean (SEM).

## ACKNOWLEDGMENTS

We thank Joanna Viguie, Karin Jonckers and Jonathan Royaert for technical assistance. We are grateful to Annette Gärtner, Eleonora Rosina, Laura D’Andrea, Esperanza Fernández, and Vittoria Mariano for helping during the development of this project and for sharing preliminary data. We would like to thank Graham Knott for their assistance in imaging and image processing, and Wim Vanduffel (KU Leuven) for supporting Marcelo Armendáriz. We are grateful to Muna Hilal for helping in setting up the MEA system and to Leonardo Restivo (NEUROBAU) for behavior advice. This work was supported by KU Leuven funds (OTF), FWO G088415N, NCCR Synapsy 51NF40-158776 and Novartis. N.D.I. is recipient of an FWO aspirant fellowship. D.S. is recipient of an FWO (12R1119N) and IWT (13160) fellowships.

## AUTHOR CONTRIBUTIONS

Conceptualization, N.D.I. and C.B.; N.D.I. participated in all the experiments described in this work. MRI imaging, D.S. and A.V.d.L., electrophysiology, A.V., M.T. and M.M, Custom-written R scripts for MEA and behavior analysis, A.C.L., Custom-made MATLAB code for EM analysis and DTI maps, M.A.; V.M, D.G., T.A., K.W.L. and A.B.S, helped with data analysis and manuscript preparation.

